# Transplanting as a means to enhance crop security of fodder beet

**DOI:** 10.1101/056408

**Authors:** EN Khaembah, WR Nelson

## Abstract

Fodder beet has become a popular winter feed for all stock classes in New Zealand. However, poor crop establishment frequently leads to either significant loss in yield, through below-target plant populations, weed competition, or crop failure. This study demonstrates that establishing the crop from transplants, common in the vegetable industry, is one way to achieve a uniform plant population and reduce weed competition through early establishment of canopy cover. The most significant effect of transplant establishment is that the target plant population is readily achieved.

## Introduction

In New Zealand, increasing interest in fodder beet (*Beta vulgaris* L.) as a winter and spring/autumn shoulder feed for ruminants has resulted in a rapid expansion of the area sown in this crop. In 2006 the cropped area was only 100 ha but this surged to an estimated 16,000 ha in 2013 (Gibbs 2014). Current industry indications are that sowing in 2015 was about 60,000 ha. Interest in the crop is driven by its high dry matter yield (DMY), high energy ratio (ME ≥12 MJ/kg DM), high palatability and flexibility in feeding (Gibbs 2014; Milne et al. 2014). Fodder beet is therefore a crop of increasing importance in New Zealand and is set to complement forage brassicas (e.g. forage kale and swedes) commonly used as winter supplementary feed.

The DMY potential of this crop is reputed to be as high as 25–28 t/ha (Matthew et al. 2011), but common commercial crops are generally less than 20 t/ha (Scott & Maley 2010). A recent evaluation of 11 cultivars marketed in New Zealand indicated DMY of 13.5–23 t/ha for irrigated and 12.7–21.9 t/ha for rain-fed crops (Milne et al. 2014). The crop is usually precision drilled using pelleted seed, but this method of establishment often results in below-target plant populations and uneven stand establishment. Drilling early in spring aims to maximise yield potential through the longer period available to intercept radiation, but sowing into cool soils frequently results in slow and erratic crop emergence. Fodder beet crops are slow to achieve canopy closure, which, combined with patchy establishment, predisposes the crop to weed competition, severely reduces yields and increases the risk of crop failure. Another constraint imposed by slow crop establishment is that the canopy closes after the radiation maximum of the year, further penalising yield.

Seed priming techniques (Jalali & Salehi 2013) may relieve some of these constraints, but this is not currently available for New Zealand fodder beet crops where seed is imported already pelleted and thus not suited to further priming techniques. It is therefore suggested that transplanting can be used as an alternative method to ensure rapid stand establishment, early season weed control, a long growing season and enhanced field-scale yields of fodder beet crops.

Transplanting techniques have been widely researched in sugar beet, an economically important crop and a close relative of fodder beet, both being of the species *Beta vulgaris*. Research trials have particularly focused on increasing sugar yield potential, as this crop is strongly responsive to length of growing season (Dillon et al. 1972; Theurer & Doney 1980; Hussain & Field 1991; Karbalaei et al. 2012), and have also focussed on weed management (Kouwenhoven et al. 1991). Although increased yield was a common factor in these trials, the value of the increase in yield compared with the increased cost of these early transplant production systems has not justified conventional sugar beet production to change to establishment via transplants. We report here early stage observations from an unreplicated proof-of-concept case study of fodder beet established from cell-grown seedlings transplanted at two rates, and the conventional precision-drilling method.

## Methods and Materials

The trial was conducted at the Lincoln research site of The New Zealand Institute for Plant & Food Research Limited. Seedlings of a commercial fodder beet (‘Rivage’, Agricom Ltd, Christchurch) were glasshouse-raised in model 144 Transplant Systems cell trays (Fig. 1) and transplanted by hand on 06 November 2014 when they were 6 weeks old. Two transplanting rates: 50 cm inter-row by 30 cm intra-row (“Transplanted (50x30)”) and 30 cm inter-row by 30 cm intra-row (“Transplanted (30x30)”), were evaluated. The crop was established alongside a replicated seed-sown trial precision-drilled at 110,000 seeds/ha (inter-row spacing = 50 cm) on 17 October 2014. Pre-sowing/transplanting fertiliser and topdressings were applied as per common agronomic recommendations (e.g. Chakwizira et al. 2014). Irrigation and herbicides were applied as required. A portable Sunfleck ceptometer (AccuPAR model PAR-80; Decagon, Pullman, WA, USA) was used to measure canopy interception approximately fortnightly during crop growth. On 16 June 2014, plants in two inner-rows over 4 m length (transplants) and two rows over 2 m length per plot (precision drilled) were harvested, counted and weighed to determine the total fresh weight. Sub-samples of two plants from each sample were retained for DM weight determination. These were washed to remove soil, re-weighed, cut into small pieces and oven-dried at 60°C until constant weight for 24 h to determine DM yield.

**Figure 1:**
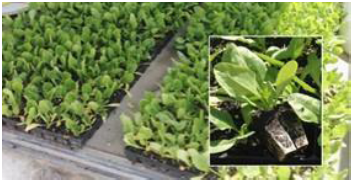
Thirty-eight-day-old fodder beet (cultivar ‘Rivage’) seedlings in model 144 Transplant Systems cell trays.

## Results and Discussion

All transplants survived, resulting in uniform stands and achievement of the target plant populations. By contrast, the drilled crops were uneven (Fig. 2) and the plant population varied by ∼10% at various points across plots (Table 1). Transplanted plots had visually lower weed infestation and required one less herbicide application. The “bulbs” of the transplanted crops had a degree of secondary root proliferation, i.e. sprangling (Fig. 3) and were considerably easier to pull from the ground. Dry matter yield was similar for the three treatments (Table 1, Fig. 4). This suggests that the advantage of earlier canopy closure of the transplanted over the precision-drilled crops (Fig. 5) did not noticeably translate into higher yield. Plant populations were in increments of ∼20,000 plants/ha across the three treatments and per plant reflected an expected trend common to different plant populations regardless of establishment method (Fig. 4). Thus, from these data it is not possible to separate yield effects from uneven crop establishment, earlier canopy cover or establishment method.

**Figure 2:**
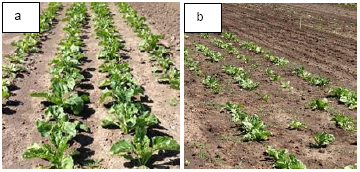
Transplanted (a) and precision drilled (b) fodder beet (cultivar ‘Rivage’) plots at the New Zealand Institute for Plant & Food Research, Lincoln, in 2014.

**Figure 3:**
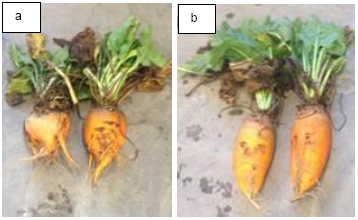
Transplanted (a) and Precision-drilled (b) fodder beet (‘Rivage’) plants at harvest. The forked root system (sprangling) of the transplanted crop is obvious and in spite of an apparent visual volume difference dry matter yields were similar.

**Figure 4:**
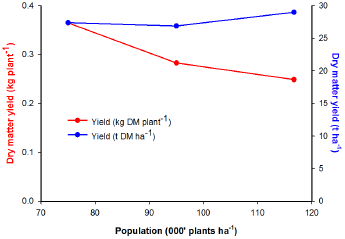
Calculated per plant and per ha dry matter yield for observed plant populations of transplanted and precision-drilled fodder beet crops.

**Figure 5:**
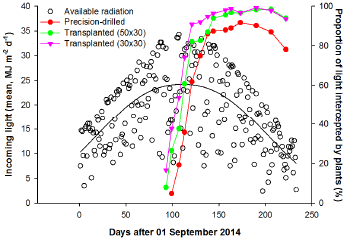
Available and intercepted radiation by transplanted and precision-drilled fodder beet crops.

**Table 1:** Plant population, per ha and per plant yield data of transplanted and precision-drilled fodder beet crops. Data for the precision-drilled are means of four replicates (± SD).

| Establishment method and spacing | Population (plants/ha) | Dry matter yield |  |
| --- | --- | --- | --- |
|  |  | t/ha | kg/plant |
| Transplanted (50x30) | 75,000 | 27.4 | 0.365 |
| Transplanted (30x30) | 116,667 | 29.0 | 0.249 |
| Precision-drilled | 95,000 (9,600) | 26.9 (2.68) | 0.283 |

It is important to note that the precision-drill rate of 110,000 seeds/ha to achieve the average 95,000 plant/ha in this experiment was higher than what farmers use in their fields (i.e. drill 100,000 seeds/ha to achieve 80,000 plants/ha). Thus, while a high plant population and yield comparable to the transplanted crops was achieved in this experiment, it was far from the 15.9 t/ha observed in a recent trial (Milne et al. 2014).

The extra speed of growth and higher potential yield, particularly as indicated by intercepted radiation (Fig. 5), is commonly reported in sugar beet transplant trials. The most likely yield increase arises from both quicker leaf canopy closure, reduced weed pressure and particularly the potential to raise seedlings earlier than is feasible by direct seed-sowing in the field. In this proof of concept trial we did not attempt to exploit this earlier plant germination component offered by transplanting.

Sprangling of roots is commonly noted in sugar beet transplant trials (Dillon et al. 1972; Theurer & Doney 1980). Severe sprangling, associated with the age of seedlings at transplanting, has been reported to reduce total yield (Dillon et al. 1972). A point of significance is that the seedlings used in these earlier transplant trials were largely bare-root as opposed to most modern systems where the entire root ball is transplanted. Bare-rooted plants are particularly susceptible to damage during handling. Sprangling was noted in the transplanted roots in this experiment, and photographic comparisons indicate the degree of sprangling was similar to that observed in sugar beet, which had no effect on yield (Theurer & Doney 1980).

Although commonly referred to as a bulb, the storage organ of sugar and fodder beet is primarily a swollen tap root. Technically, the swollen storage organ is comprised of a shallow crown portion with leaves attached, followed by the neck as the widest proportion derived from the hypocotyl, and the remainder of the depth comprising the primary tap root (Artschwager 1926). This mild sprangling is therefore the result of growth of secondary roots following pruning to the tap root apex. This will occur in 100% of transplanted seedlings incorporating root pruning at the base of the container. Sprangling has not generally been noted as an issue associated with reduced yield in sugar beet, and this trial suggests the mild sprangling noted in fodder beet transplants has not impacted yield either. However, although a completely subjective measure, the ease of lifting mildly sprangled roots may be of significance where crops are grown for lifting, or possibly even ease of access to the “bulb” by the grazing animal.

Transplanting of cell-grown seedlings is a well-established technique used worldwide for establishment of many vegetable crops. The relatively small New Zealand population and lack of export opportunities for fresh vegetables has resulted in a relatively small-scale nursery system and most vegetable seedlings are transplanted manually or with various common, mechanically robust, semi-automatic field transplanters. The processing tomato crops grown in New Zealand’s Hawke’s Bay region are more analogous in terms of scale and short window for transplanting. A much higher degree of automation is evident within this industry, including automated field transplanting at high speed.

Should the animal-feed industry choose to develop fodder beet as a transplanted crop, two further areas for refinement of the technique suggest themselves. Firstly, early crop establishment is critical to increasing yield potential. At present this is largely limited by soil temperature suitable for seed germination and thus crop establishment typically in the September/October period. Protected and heated growth conditions in a nursery make early production of seedlings possible and therefore the window of opportunity for field transplanting can be brought forward depending on land access. Secondly, fodder beet varieties are very different in their growth habits, with some “bulbs” produced largely underground while others are largely above ground. If sprangling proves to be detrimental to total consumable yield, then a focus on varieties where storage is more in the neck (hypocotyl) portion of the storage organ might prove beneficial.

## Conclusion

This study demonstrates that transplanting technique can be used to achieve target plant populations of fodder beet crops, reduce the use of herbicides and capitalise on a longer effective growing season through early canopy closure. Further research refinements on the cell transplant technique is required to enable full capture of the benefits of earlier canopy closure and superior light interception in transplanted crops. Technology within nurseries and for field transplanting is available internationally and therefore scaling up will not require new technology, but simply development of businesses to offer these services on the scale required to service the fodder beet industry.

## Acknowledgments

The authors thank Shane Maley and Mike George for their contribution to the trial. Funding for the transplanting project was provided by The New Zealand Institute for Plant & Food Research Limited core funds (Blue Skies #413). The precision-drilled experiment was funded by the Forages for Reduced Nitrate Leaching programme with principal funding from the New Zealand Ministry of Business, Innovation and Employment. The programme is a partnership between DairyNZ, AgResearch, Plant & Food Research, Lincoln University, Foundation for Arable Research and Landcare Research.

